# Multimodal neuroimaging–microbiota integration identifies *Akkermansia* as a modulator of alcohol-induced gut–liver–brain pathology

**DOI:** 10.64898/2026.08.03.742281

**Authors:** Mohamed Kotb Selim, Daniel Panadero Soler, Silvia De Santis, Alfonso Benítez-Páez, Alejandra Flor, Carlos Sanz, Mariana Mesquita, Francisco Javier Cubero, Roberto Ciccociopo, Antonio Pertusa, Yolanda Sanz, Santiago Canals

## Abstract

Alcohol use disorder (AUD) disrupts the gut–liver–brain axis, yet mechanistically grounded and therapeutically actionable targets within this network remain poorly defined. To identify microbial modulators of alcohol-induced tissue pathology, longitudinal advanced diffusion MRI and fecal 16S rRNA profiling were integrated across Marchigian Sardinian alcohol-preferring rats evaluated at baseline, after four weeks of voluntary alcohol intake, and following six weeks of abstinence. Machine learning, specifically random forest models combining neuroimaging and microbiota data, improved phase classification and identified Akkermansia as the microbial feature most strongly associated with alcohol-related white matter microstructural abnormalities. Alcohol exposure induced widespread white matter alterations alongside gut dysbiosis characterized by reduced microbial diversity. To evaluate functional relevance, Akkermansia muciniphila was administered during the abstinence phase. Supplementation with A. muciniphila restored intestinal mucus, reduced liver injury markers, and elevated myelin basic protein levels within affected white matter regions. Collectively, these findings highlight Akkermansia as a critical modulator of alcohol-induced gut–liver–brain pathology and provide experimental support for a causal contribution of specific gut bacteria to persistent white matter damage in AUD. More broadly, this work establishes a robust multimodal framework for microbiome-based target discovery with clear translational relevance for disorders characterized by dysfunction along the gut–liver–brain axis.

**Research in context:** *Evidence before this study:* Alcohol use disorder (AUD) is associated with gut dysbiosis, impaired intestinal barrier function, liver injury, and persistent white matter abnormalities. Previous studies in patients and animal models have linked alcohol exposure to reduced microbial diversity, altered gut permeability, and white matter microstructural damage, particularly during abstinence. Other work has shown that microbiota-derived interventions can ameliorate peripheral consequences of alcohol exposure, especially in the gut and liver. However, the specific microbial features linked to alcohol-induced brain pathology remain poorly defined, and no prior study has integrated longitudinal microbiota and neuroimaging data to identify candidate microbial modulators of alcohol-related white matter damage and then functionally test them in vivo across the gut–liver–brain axis.

*Added value of this study:* We developed a multimodal framework that integrates longitudinal advanced diffusion MRI with fecal microbiota profiling and machine learning in alcohol-preferring rats. This approach identified *Akkermansia* as the microbial feature most strongly associated with alcohol-induced white matter abnormalities. Guided by this result, we administered *Akkermansia muciniphila* during abstinence and observed coordinated beneficial effects across multiple organs, including restoration of intestinal mucus, reduction of liver injury markers, and recovery of myelin basic protein in affected white matter regions. To our knowledge, this is the first study to combine longitudinal microbiota–MRI integration with experimental validation of a microbiota-based intervention that mitigates alcohol-induced pathology across the gut–liver–brain axis while restoring central white matter integrity.

*Implications of all the available evidence:* Our findings support a mechanistic contribution of specific gut bacteria to persistent alcohol-induced tissue damage and identify *Akkermansia* as a candidate modulator of gut–liver–brain axis dysfunction in AUD. More broadly, this study establishes a generalizable strategy for integrating microbiota and neuroimaging data to discover biologically meaningful and therapeutically actionable targets in complex disorders involving coordinated peripheral and central pathology.

## INTRODUCTION

Alcohol use disorder (AUD) is a chronic relapsing condition characterized by persistent alcohol consumption despite adverse consequences, leading to substantial impairments in health and daily functioning^1^. It represents a major global health challenge: in 2019, more than 400 million individuals worldwide were living with AUD, with alcohol consumption accounting for approximately 2.6 million deaths^2^. Accordingly, alcohol use contributes 5.1% of disability-adjusted life years and has been causally linked to more than 200 diseases and injury conditions, including liver disease, cancer, and cardiovascular disorders^3^.

Chronic alcohol consumption leads to widespread and persistent pathological changes throughout the body^4,5^. In the brain, sustained alcohol exposure causes microstructural damage to white matter, particularly in the corpus callosum, fimbria, and fornix, with these alterations persisting into early abstinence^6,7^. Peripherally, alcohol exerts multiple effects, including impairment of intestinal barrier integrity and alterations in gut microbiota composition and function, a phenomenon known as dysbiosis^8,9^. These organ-specific effects are interconnected, as chronic alcohol use disrupts the coordinated interactions between physiological systems^10^, with inflammation emerging as a central pathological mechanism^4^. Notably, both clinical and preclinical evidence suggests that changes in the gut microbiota may occur before overt systemic manifestations of AUD^4,11^.

This integrative perspective is supported by accumulating evidence indicating a causal role of the gut microbiota in modulating brain function^12^. For instance, administration of specific probiotic species has been shown to modulate central neurotransmitter systems, such as GABAergic and dopaminergic pathways^11,13,14^, which are critically involved in addiction-related behaviors. Despite these advances, the precise microbial changes associated with alcohol-induced brain damage remain poorly characterized. This knowledge gap underscores the need for multimodal and longitudinal approaches to identify microbiota signatures that are specifically linked to alcohol-related neural alterations.

In this study, we hypothesized that integrating gut microbiota profiles with brain MRI measures would reveal microbiota changes specifically linked to alcohol-induced neural damage. To test this hypothesis, we performed longitudinal fecal microbiota profiling and diffusion MRI-based white matter analyses in genetically selected alcohol-preferring Marchigian sardinian (msP) rats, an established model for excessive alcohol drinking^15^. Assessments were conducted at baseline, during alcohol exposure, and throughout abstinence. We then applied a data-driven machine-learning framework to integrate microbiota and neuroimaging features, aiming to identify microbial shifts predictive of white matter alterations. Finally, we tested the functional relevance of these findings by administering the identified candidate bacterial species to alcohol-exposed rats and evaluating its effects on the gut, liver, and brain. Collectively, our study demonstrates a multimodal approach to uncover gut–brain axis interactions in AUD and suggests that targeting the microbiota may offer a promising therapeutic strategy.

## METHODS AND MATERIALS

### Animals

The Marchigian sardinian genetically selected alcohol-preferring (msP) rat line was used, as these animals have been selectively bred for high voluntary alcohol consumption and preference^14^. In addition, msP rats display anxiety-like behaviors and heightened stress sensitivity, making them a well-established model for investigating the neurobiological mechanisms underlying excessive alcohol consumption and potential intervention for AUD.

msP rats were obtained from the host breeding facility at the School of Pharmacy, University of Camerino in Camerino Italy. Upon arrival to the Instituto de Neurociencias in Alicante, animals were group-housed and habituated for five days, after which they were individually housed with environmental enrichment. Housing conditions were maintained at 22 ± 2 °C and 55 ± 10% humidity under a 12-hour light–dark cycle, with ad libitum access to food and water.

For alcohol exposure, rats were subjected to a two-bottle free-choice paradigm, consisting of water and a 10% (v/v) ethanol solution (absolute ethanol, Panreac Química SLU) for four weeks. Fluids were refreshed regularly, and bottle positions were alternated to prevent side preference. Body weight and fluid intake were recorded every 2–3 days.

### Study Design

We conducted three experiments: one longitudinal MRI study with fecal sampling and two cross-sectional histological studies. Animals were randomly assigned to experimental groups, and investigators were blinded to group allocation during data acquisition and analysis.

In Experiment 1, a total of 27 male Marchigian Sardinian alcohol-preferring (msP) rats (370–480 g, 8 weeks old) were used (Fig. 1a). Animals were allocated to two groups: (i) an MRI cohort undergoing the two-bottle free-choice alcohol paradigm (n = 18) and (ii) an MRI control cohort receiving only water (n = 9). In the MRI cohort, longitudinal imaging, together with fecal sample collection, was performed across three experimental phases: baseline (TP1), after four weeks of alcohol exposure (TP2), and following six weeks of abstinence (TP3). These longitudinal MRI data from the alcohol-exposed group were used for subsequent machine learning analyses together with the microbiota analysis performed at the same TPs. The water-drinking control group was included for comparison with baseline and alcohol-exposed conditions but was not incorporated into the machine learning models.

**Figure 1:**
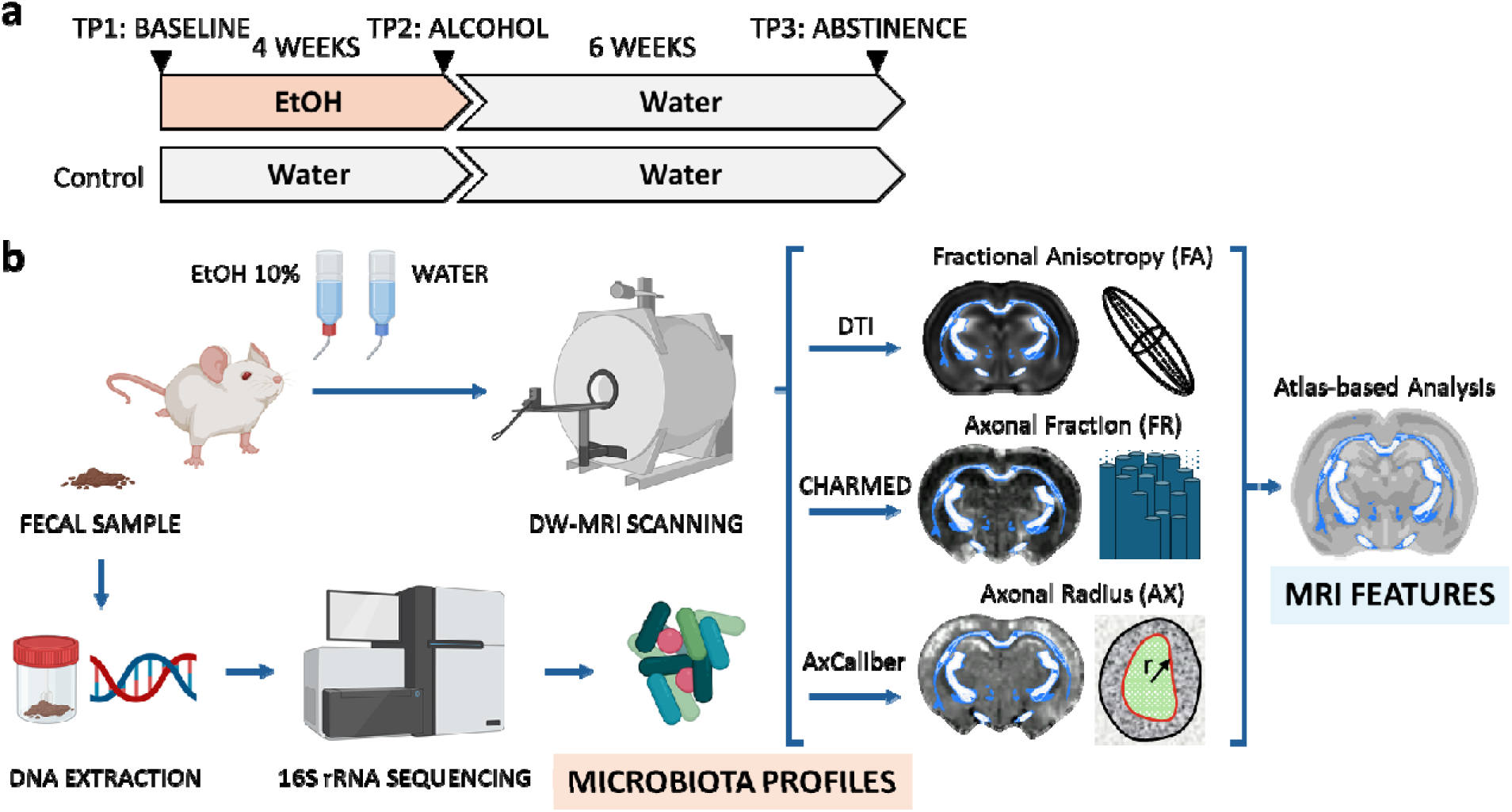
Study design. (a) Experimental design and timeline of data acquisition across the different study phases. (b) Schematic overview of the experimental workflow, including data collection at each longitudinal timepoint (TP1–TP3) and extracted features.

Experiment 2 was designed to investigate the histological correlates of alcohol-induced white matter alterations identified in Experiment 1. An independent cohort of male msP rats (n = 10; Fig. 3a) was used for immunohistochemical analyses following alcohol exposure.

Experiment 3 was designed to evaluate the effects of *Akkermansia muciniphila* supplementation during abstinence and included a total of 36 msP rats, with equal representation of males and females. This experiment followed the same temporal structure as Experiment 1, with bacterial supplementation administered throughout the abstinence period. Two water-drinking control groups were included, one of them receiving the same bacterial supplementation as the corresponding alcohol drinking group (Fig. 6a).

Experiments 1 and 2 included only male rats, whereas Experiment 3 included both sexes to assess generalizability.

### Fecal Sample Collection, DNA extraction and library construction

Fecal samples were collected by placing the animal in a sterile cage and gently massaging their abdomen to stimulate defecation. Around two to three freshly defecated pieces were collected using sterile forceps and placed into a 3 mL Eppendorf tube. The tubes were then transferred to dry ice before storage at −80 °C until needed for microbiota profiling.

DNA was extracted from rat fecal samples using the QIAamp® PowerFecal® DNA Kit (Qiagen, Germany) according to the manufacturer’s instructions. Cell lysis was carried out using a Mini-Bead Beater apparatus (BioSpec Products, USA) and DNA concentration was measured using a Nanodrop (Thermo Fisher Scientific, USA). The V3-V4 hypervariable regions of the 16S ribosomal ribonucleic acid (rRNA) gene were amplified using ∼10 ng DNA and 25 PCR cycles consisting of the following steps: 95 °C for 20 sec., 55 °C for 20 sec. and 72 °C for 20 sec. Phusion High-Fidelity Taq Polymerase (Thermo Fisher Scientific, USA) and the 6-mer barcoded primers, S-D-Bact-0341-b-S-17 (TAGCCTACGGGNGGCWGCAG) and S-D-Bact-0785-a-A-21 (ACTGACTACHVGGGTATCTAATCC)^44^ were used for PCR. Dual-barcoded PCR products were purified from triplicate reactions using the Illustra GFX PCR DNA and Gel Band Purification Kit (GE Healthcare, UK) and quantified using a Qubit 3.0 fluorometer with the Qubit dsDNA HS Assay Kit (Thermo Fisher Scientific, USA). Multiplexing was carried out by combining equimolar quantities of V3-V4 amplicons (∼50 ng per sample) and sequenced on one lane of Illumina MiSeq platform with 2×300 PE configuration (Centro Nacional de Analisis Genómico - CNAG, Barcelona, Spain).

### Image Acquisition and Analysis

Imaging experiments were carried out under isoflurane anesthesia, which was induced using 4-5% isoflurane in oxygen (0.8-1L/min). Animals were held in a custom-made holding apparatus featuring a tooth bar and a nose cone. Throughout scanning, isoflurane concentration was maintained at 1.2%, while body temperature was monitored and kept constant at 37± 0.5 °C using a heating pad. The physiological parameters, including oxygen saturation, pulse, and breathing rate, were monitored (MouseOx, Starr Life Sciences, Oakmont, PA, USA).

Magnetic Resonance Imaging (MRI) was performed on a horizontal 7 T scanner with a 30 cm diameter bore (Biospec 70/30, Bruker Medical, Ettlingen, Germany) at the Institute of Neurosciences in Alicante (CSIC-UMH). We used a receive-only phase array coil with an integrated combiner and preamplifier, combined with an actively detuned transmit-only resonator. Diffusion-weighted data was acquired using an EPI spin-echo diffusion sequence. We scanned a protocol with 30 uniformly distributed gradient directions, b-value = 1000 and 2500 s/mm², with three non-diffusion weighted images (b0), repetition time (TR) = 8000 ms, and echo time (TE) = 29 ms. Sixteen coronal slices were imaged for each subject (field of view [FOV] = 32 × 32 mm², matrix size = 110 × 110 × 16, in-plane resolution = 0.225 × 0.225 mm², slice thickness = 1 mm). The same protocol was repeated, using a stimulated-echo sequence (Frahm et al., 1985) at four diffusion times: 15, 30, 60, and 100 ms. T1 and T2-weighted maps were also acquired using the same settings.

Raw data was converted from the Bruker Paravision format to NIfTI (The Neuroimaging Informatics Technology Initiative) format, then reoriented and augmented by multiplying the voxel size by ten. Skull stripping was carried out as previously described^45^, we used the buildtemplateparallel.sh script implemented in Advanced Normalization Tools (ANTs)^46^. Registration was used to mask the brain in native space, and each sample was manually checked and corrected to ensure each brain had been successfully extracted to create a binary mask as required for downstream processing.

Fractional Anisotropy (FA) maps were generated by first pre-processing according to community standards^47^. Steps included denoising, Eddy current, motion distortions, and bias field corrections. Diffusion Tensor Imaging (DTI) fitting was performed using a Robust Estimation of Tensors by Outlier Rejection (RESTORE) approach, which iteratively reweighs least-squares regression to identify and exclude outliers from the fit^48^.

To measure Axonal Fraction (FR), a surrogate for axonal density, pre-processed multi-shell acquisition data with b-values between 1000 and 2500 were processed through the CHARMED model^49^ using an in-house MATLAB script (The Mathworks Inc., Natick, MA).

Diffusion data with multiple diffusion times acquired on a Stimulated Echo Acquisition Mode (STEAM) were pre-processed by combining features of a custom made pipeline using the Nipype package^50^ with tools from the DIPY package^51^. This pipeline was made publicly available on Github (https://github.com/mokselim/Preprocessing_msP_diffusion) (**Supp. Fig. 5**) and provides a measure of Axonal Diameter (AX).

For ROI-based analyses, each animal’s FA map was compared from TP2 to itself at TP1 to reduce non-biological variance of the same brain imaged at two different times. TP1 maps were next registered nonlinearly to the SIGMA rat template^52^. Both registrations were applied to the FR and AX maps using the antsApplyTransforms command in the ANTs package. ROI’s for the corpus callosum, fimbria and fornix from the SIGMA atlas were used to mask data to extract the mean value per ROI.

All three measures, Fractional Anisotropy (FA), Axonal Fraction (FR), and Axonal Diameter (AX), for both alcohol exposed and control animals, underwent whole-brain statistical analysis. A modified version of tract-based spatial statistics (TBSS)^53^ was used in combination with an advanced normalization approach for adaptation to animal data. A general linear model (GLM) was used within a voxel-wise, non-parametric permutation-based statistical framework to test for significant differences between the groups. Threshold Free Cluster Enhancement was used to account for multiple comparisons across clusters. We performed 10,000 permutations, and a voxel-wise p-value corrected for multiple comparisons was considered statistically significant if it was less than 0.05. Cluster locations were achieved using a white matter FA-driven skeleton by averaging all subjects using the tbss_skeleton routine in FSL (FMRIB Software Library)^54^.

### Immunohistochemistry, histological evaluation and cytokine measurement

Animals in the immunohistochemistry group were injected intraperitoneally with a lethal dose of sodium pentobarbital, 46 mg/kg (E.V.S.A. laboratories, Madrid, Spain), then perfused intracardially with 100 ml of 0.9% phosphate saline buffer (PBS) and 100 ml of ice-cold 4% paraformaldehyde (PFA, BDH, Prolabo, VWR International, Louvain, Belgium).

The brains were next removed from the skull and fixed in 4% PFA for 1 hour, then embedded in 3% agarose/PBS (Sigma-Aldrich, Madrid, Spain) and cut into 50-μm-thick serial coronal sections using a vibratome (VT 1000 S, Leica, Wetzlar, Germany). Coronal sections underwent three rinses in 1 x PBS containing 0.5% Triton X-100 (Sigma-Aldrich, Madrid, Spain) for 10 minutes each, followed by a 2-hour blockade in the same solution with 4% bovine serum albumin (Sigma-Aldrich, Madrid, Spain) and 2% goat serum (Sigma-Aldrich) at room temperature. Slices were incubated overnight at 4 °C with primary antibodies against neurofilament 160 kD medium (1:250, Abcam Cat# ab134458) and myelin basic protein (1:250, Millipore Cat# MAB384-1ML) to label axonal processes and myelin, respectively. The next day, secondary antibodies conjugated to fluorescent probes (Molecular Probes Cat# A-11029; Molecular Probes Cat# A-11042) were added to the sections at a concentration of 1:500 and incubated for 2 hours at room temperature. The sections were then treated with 15 mM 4′,6-Diamidine-2′-phenylindole dihydrochloride (DAPI, Sigma-Aldrich) for 15 minutes at room temperature. Subsequently, the sections were mounted on slides and covered with an anti-fading medium made from a 1:10 solution comprising Propyl-gallate and Mowiol (P3130, Sigma-Aldrich; 475904, MERCK-Millipore, Massachusetts, United States). Note, for myelin labelling, antigen retrieval was performed using 1% citrate buffer (Sigma-Aldrich) and 0.05% Tween 20 (Sigma-Aldrich) warmed to 80 °C to unmask its proteins.

Slides were then analyzed using a computer-assisted morphometry system that included a Leica DM4000 fluorescence microscope equipped with a QICAM Qimaging camera (model 22577, Biocompare, San Francisco, USA) and Neurolucida morphometric software (MBF, Biosciences, VT, USA). Myelin and neurofilament fluorescent analyses were performed using Icy software^55^, and ImageJ^56^. Two rectangular ROIs of 400×300 µm2 each were placed, one for the midsection of corpus callosum, and one for fimbria-fornix bundle per hemisphere in at least 2 slices per rat to obtain the corresponding intensity values.

4% Formalin-fixed paraffin-embedded sections of liver and intestine were de-paraffinized and stained with hematoxylin and eosin (H&E), mucin 2 (sc-515032, Santa Cruz) and adipophylin perilipin (Ab108323, Abcam). Nuclei were counter-stained with hematoxylin. Tissues were coded at time of collection to assure an un-biased analysis; at least 3 images were acquired per tissue section (Bright Field Microscope DM1000, Leica, Wetzlar and ZEISS Axio Lab A1. Carl Zeiss Microscopy GmbH, Jena) and semi-quantification of positive staining was performed using Image J software (http://imagej.nih.goc/ij/, National Institutes of Health, Bethesda, MD). Histological scores were performed by an experienced pathologist. The concentration of TGFß was measured following the manufactureŕs protocol (R&D, Ref. DB100C).

### Microbiota Analysis

Raw data was received in the form of fastq files, pair ends with quality filtering were assembled using Flash software^57^. Assessment of alpha and beta diversity analyses was performed using the Operational Taxonomic Unit (OTU)-picking approach. In brief, after de-multiplexing and barcode/primer removal, chimera detection was performed using the UCHIME algorithm^58^ and the SILVA seed reference set of 16S rRNA sequences (Release 138)^59^. OTU information was retrieved by using a rarefield subset of 10,000 sequences per sample, which were randomly selected after multiple shuffling (10,000X) from the original dataset, and the UCLUST algorithm implemented in USEARCH v8.0.1623^60^.

Alpha diversity descriptors, Chao’s index, Simpson’s reciprocal index, Simpson’s evenness and dominance index were calculated using QIIME v1.9.1^61^.

The community structure across the sample groups was determined using the vegan v2.6.4 R package through exploratory multivariate appraisal based on the non-metric multidimensional scaling (NMDS) (vegan::metaMDS R function and “bray” distance).

For taxonomy assessment, the complete set of sequences obtained after chimera removal was used. Following the OTU-picking approach (UCLUST algorithm), the transformation of compositional microbiota-derived data was achieved by applying the centered log-ratio (clr) algorithm implemented in *CoDaSeq::codaSeq.clr* R function with prior execution of *zCompositions::cmultRepl* function supporting a Bayesian-Multiplicative replacement of count zeros. Taxonomy identification for OTUs of interest was assisted by utilization of SINA aligner^62^ and SILVA database (Release 138).

Pairwise t-test or Wilcoxon Rank Sum test were used to test differences in time points for comparisons across the alpha diversity descriptors, with prior evaluation of normal distribution of variables (Shapiro-Wilk test). Differences in the microbial community structure were assessed by interpretative approaches using the permutation-based *vegan::adonis2* function. Differential abundance of OTUs across time points was also assisted by application of LMM (linear mixed models) controlling idiosyncratic variation across animal specimens. Graphs and plots were drawn using the ggplot2 R package (R distribution v4.4).

### Culture conditions of Akkermansia muciniphila and administration to rats

### Akkermansia muciniphila

DSM 22959 was grown in Medium 104 supplemented with 0.1% mucin at pH 7.2, as recommended by the German Collection of Microorganisms and Cell Cultures (DSMZ). To sterilise the mucin before adding it to the medium, it was covered with 95% ethanol and incubated at 70 °C for 24 h, allowing the alcohol to evaporate and then the mucin was added to the medium. Agar plates were introduced into the anaerobic chamber 1 day prior to use to remove oxygen from the medium. *A. muciniphila* DSM 22959 was inoculated in agar plates of the recommended medium and incubated in an anaerobic chamber (10% CO2, 10% H2, 80% N2) at 37°C for 4 days. Subsequently, the cell biomass was collected from the agar surface using a swab and suspended in PBS containing 0.05 % cysteine and 20% glycerol. Aliquots were frozen in liquid nitrogen, checked by sequencing the 16S rRNA gene and kept at −80°C until use. The bacterial strain concentration was quantified by plate counting, and 10^8^ CFU/day were administered to each animal.

To facilitate voluntary oral intake of the bacterial strain, the experimental dose was incorporated into a flavored gelatin matrix. To prepare the mixture, 60 mL of water and a gelatin sheet were microwaved until the sheet fully dissolved or the water reached boiling. Upon removal from the microwave, 5g of strawberry-flavored gelatin powder was added and stirred until completely dissolved, followed by the addition of 60 mL of water and thorough mixing. The resulting solution was poured into molds and refrigerated at 4°C for 3–4 hours (preferably overnight) to allow complete gelation. The control group received the same gelatin preparation without the addition of *A. muciniphila*.

Using a sterilized spoon, approximately 1–1.5 g of gelatin was extracted per dose. Then, 100 µL of either vehicle (culture medium; control group or an *A. muciniphila* suspension (treatment group) was injected into multiple sites within the gelatin to ensure homogeneous distribution. The prepared gelatin dose was held beneath the cage grid, allowing for voluntary consumption. Rats that initially displayed hesitation or required more time to consume the gelatin (typically during the first two to three exposures) generally adapted to its consumption in subsequent administrations.

Rats were supplemented with *A. muciniphila* or vehicle via gelatin on a daily basis (weekends excluded) for six weeks. After this time period, rats were euthanized and immunohistochemistry and histological evaluation were carried out as described above in ‘Immunohistochemistry and Histological Evaluation’.

### Random Forest Modeling for Phase Prediction

Two datasets, microbiota composition and MRI-derived features, from Experiment 1 (see Study Design) were integrated following Ong et al.^63^. Each dataset comprised 54 samples (18 per phase: baseline, alcohol exposure, and abstinence), with 172 features in the microbiota dataset and 144 features in the MRI dataset. Microbiota data were log-transformed, and MRI features were z-scored prior to analysis.

Random Forest (RF) models were used for both classification and regression tasks. All analyses employed Leave-One-Out cross-validation (LOOCV), using 1600 estimators for classification and 4000 estimators for regression. For classification, microbiota and MRI datasets were used independently to predict experimental phase (baseline, alcohol exposure, or abstinence). For regression, microbiota features were used to predict individual MRI-derived metrics.

Feature relevance was assessed using permutation importance. For each feature, model performance was re-evaluated after random shuffling of its values, and the resulting decrease in classification accuracy or increase in mean squared error (for regression) was quantified. To improve robustness, importance scores were averaged across 30 independent permutations per feature.

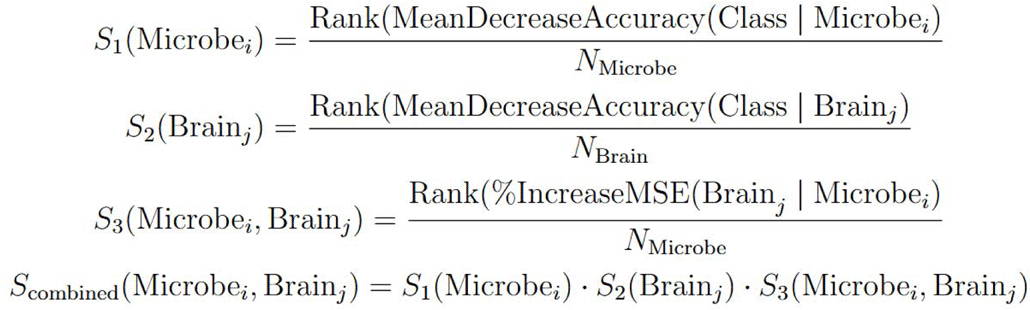

Based on this approach, three importance scores were defined. S1 and S2 represent the normalized importance rankings of microbiota and MRI features, respectively, derived from RF classification. S3 represents the normalized importance of microbiota features for predicting MRI features in the regression analyses. For S3, a separate ranking was obtained for each MRI feature, such that each score corresponds to a specific microbiota–MRI feature pair. This framework captures feature-specific gut–brain associations, allowing identification of microbiota features that differentially contribute to distinct MRI-derived measures.

To integrate these measures, S1, S2, and S3 were combined by direct multiplication to generate a composite score (S_combined), assigned to each microbiota–MRI feature pair. Higher S_combined values indicate stronger gut–brain associations contributing to accurate prediction of alcohol consumption phase. These associations were visualized using a circos plot, in which link widths are proportional to the relative magnitude of S_combined.

From a computational perspective, estimation of S1 and S2 required 31 RF classification runs per feature (30 permutations plus one reference model), for a total of 172 and 144 features in the microbiota and MRI datasets, respectively. For S3, this procedure was repeated for each MRI feature as a regression target, substantially increasing computational load. As LOOCV was applied throughout, each RF model involved 54 training and testing iterations.

## RESULTS

### msP rats exhibit increased alcohol consumption and preference

In this study we used genetically selected alcohol preferring msP rats, a well-characterized model of innate propensity to excessive alcohol drinking^15^. Following four weeks of exposure to the two-bottle free-choice paradigm (Fig. 1), msP rats exhibited a progressive increase in alcohol consumption relative to water (Supp. Fig. 1a), reaching consumption levels of 6.53±0.42 g/Kg/day. Alcohol preference emerged within three days and stabilized after eight days of exposure, consistent with previous reports^14^ (Supp. Fig. 1b).

### Alcohol induces widespread white matter microstructural alterations

Consistent with previous reports^6,7,15–17^, four weeks of alcohol consumption in msP rats induced widespread white matter microstructural alterations. Significant changes were observed in fractional anisotropy (FA), restricted water fraction (FR; CHARMED model), and axonal radius (AX; AxCaliber model), with prominent involvement of the corpus callosum (CC), fimbria, and fornix (Fig. 2). Two-way ANOVA revealed significant effects of time (TP1-3; baseline, alcohol exposure and abstinence) and region of interest (ROI) across all three metrics, along with a smaller but significant TP × ROI interaction, indicating region-specific longitudinal evolution of microstructural changes (Supp. Table 1a). These alterations extended beyond the selected tracts, as confirmed by voxel-wise tract-based spatial statistics (TBSS) analyses (Supp. Fig. 2).

**Figure 2.**
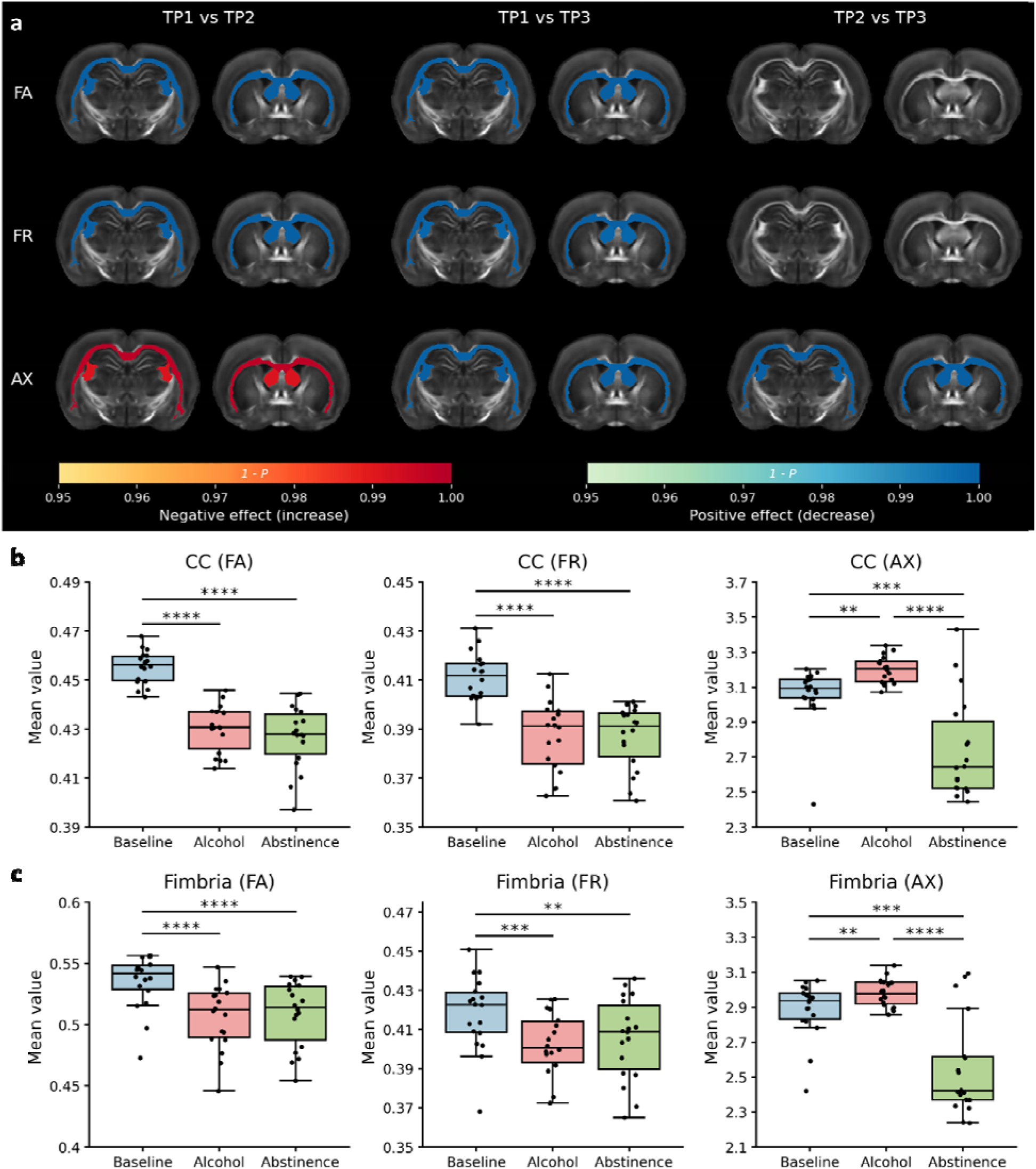
White matter microstructural alterations assessed using diffusion MRI metrics. (a) Atlas-based region-of-interest (ROI) analyses using the SIGMA rat brain template identified anatomical regions exhibiting significant changes between experimental phases: baseline (TP1), alcohol exposure (TP2), and abstinence (TP3). Color coding represents statistical significance (*p*-value), with blue indicating reductions and red increases in the corresponding metric values between timepoints. Contrasts across all three timepoints are shown for fractional anisotropy (FA), restricted fraction (FR), and axonal radius (AX), displayed on representative brain slices highlighting the corpus callosum and fimbria. (b,c) Boxplots of MRI-derived measures shown in (a) for the corpus callosum (b) and fimbria (c) across experimental phases. Boxes represent the interquartile range (25th–75th percentiles), with the median indicated by a central line; whiskers extend to 1.5× the interquartile range. Individual animals are shown as dots. Statistical significance: *p < 0.05, **p < 0.01, ***p < 0.001, ****p < 0.0001.

Post hoc statistical (paired t-test) and effect size (Cohen’s d) analyses (Fig. 2; Supp. Fig. 3) showed that alcohol consumption induced robust reductions in FA across all three regions (*CC*, t(17) = 11.1, p = 3×10⁻ □, d = 2.62; *fimbria*, t(17) = 6.67, p = 4×10⁻ □, d = 1.57; *fornix*, t(17) = 5.29, p = 6×10⁻ □, d = 1.25). Similarly, FR was significantly reduced in the *CC* (t(17) = 5.76, p = 2×10⁻ □, d = 1.36), *fimbria* (t(17) = 4.69, p = 2×10⁻□, d = 1.11), and *fornix* (t(17) = 2.62, p = 0.02, d = 0.62). In contrast, AX increased following alcohol exposure in the *CC* (t(17) = 3.65, p = 2×10⁻³, d = 0.86) and *fimbria* (t(17) = 2.94, p = 9×10⁻³, d = 0.69), but not in the fornix (t(17) = 1.57, p = 0.14).

Importantly, control animals consuming only water over the same period of time showed no significant changes in FA or FR and only a slight increase in AX (Supp. Table 2), which did not account for the alterations observed following alcohol exposure.

Post hoc statistical and effect size analyses during the abstinence period (Fig. 2; Supp. Fig. 3) showed that white matter microstructure did not recover and continued to evolve over six weeks despite cessation of alcohol exposure. Both FA and FR remained persistently reduced across all three regions, with no significant changes compared to the alcohol-exposed state (FA: *CC*, t(17) = 0.90, p = 0.38, d = 0.21; *fimbria*, t(17) = 0.43, p = 0.67, d = 0.10; *fornix*, t(17) = 0.35, p = 0.73, d = 0.08. FR: *CC*, t(17) = 0.10, p = 0.92, d = 0.02; *fimbria*, t(17) = 1.42, p = 0.17, d = 0.33; *fornix*, t(17) = 1.27, p = 0.22, d = 0.30). In contrast, AX, which had increased during alcohol exposure, showed a significant reduction during abstinence across all regions (*CC*, t(17) = 6.71, p = 4×10⁻□, d = 1.58; *fimbria*, t(17) = 6.65, p = 4×10⁻□, d = 1.57; *fornix*, t(17) = 8.03, p = 3×10⁻□, d = 1.89), indicating ongoing microstructural changes despite alcohol cessation.

### Alcohol-driven white matter alterations reflect axon–myelin unit disruption

Together, the combination of decreased FA and FR with increased AX after alcohol drinking indicates a reduction in axon–myelin unit integrity^18^. To further investigate the histological basis of these MRI findings, we analyzed an independent cohort of msP rats (n = 10; Fig. 3a). Myelin content and axonal integrity were assessed using MBP and neurofilament immunostaining, respectively (Fig. 3b). Alcohol-exposed animals showed a significant reduction in MBP staining intensity in the fimbria–fornix bundle (t(8) = 3.04, p = 0.02, d = 1.39), with no changes in control (water drinking) animals. Texture analysis revealed trends toward reduced contrast and increased homogeneity, while entropy remained unchanged (Fig. 3c).

**Figure 3.**
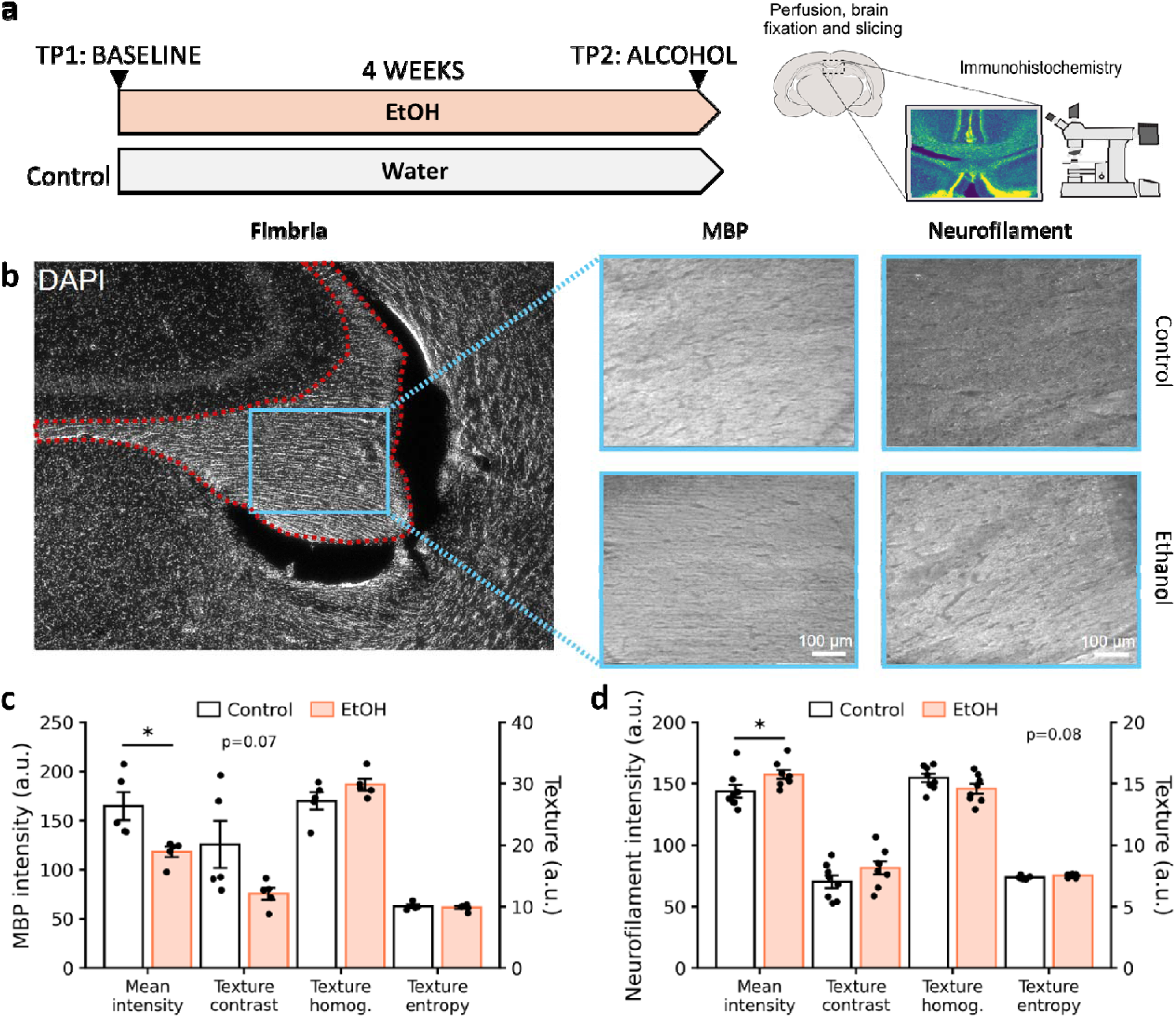
Histological assessment of white matter alterations. (a) Experimental design and timeline of data acquisition. (b) Representative coronal section stained with DAPI highlighting the fimbria bundle. (c,d) ROI-based quantitative analysis of neurofilament (c) and myelin basic protein (MBP) (d) staining intensity in the fimbria bundle. Statistical comparisons were performed using unpaired t-tests. Dashed red lines delineate the anatomical ROI, and light-blue boxes indicate representative areas used for image analysis. Statistical significance: *p < 0.05.

In contrast, neurofilament staining showed increased mean fluorescence intensity in alcohol-exposed rats (t(8) = 2.60, p = 0.03, d = 1.28), accompanied by trends toward altered texture contrast and reduced homogeneity (Fig. 3d). These findings are consistent with previous evidence indicating that diffusion MRI–derived increases in axonal diameter and reductions in axonal density reflect underlying axon–myelin pathology^18–20^

### Alcohol consumption induces gut microbiota perturbations that are largely reversed upon cessation

16S rRNA sequencing of fecal samples identified 869 operational taxonomic units (OTUs). Analysis of alpha diversity across timepoints revealed that sustained alcohol consumption reduced microbial richness (Chao index) and community evenness, consistent with increased dominance of specific taxa. Simpson’s reciprocal index further indicated a global alteration of microbial diversity during the alcohol exposure phase, which was largely restored during abstinence. Similarly, the dominance index increased during alcohol consumption and returned to baseline levels following cessation, indicating a reversible shift in community structure (Fig. 4a).

**Figure 4.**
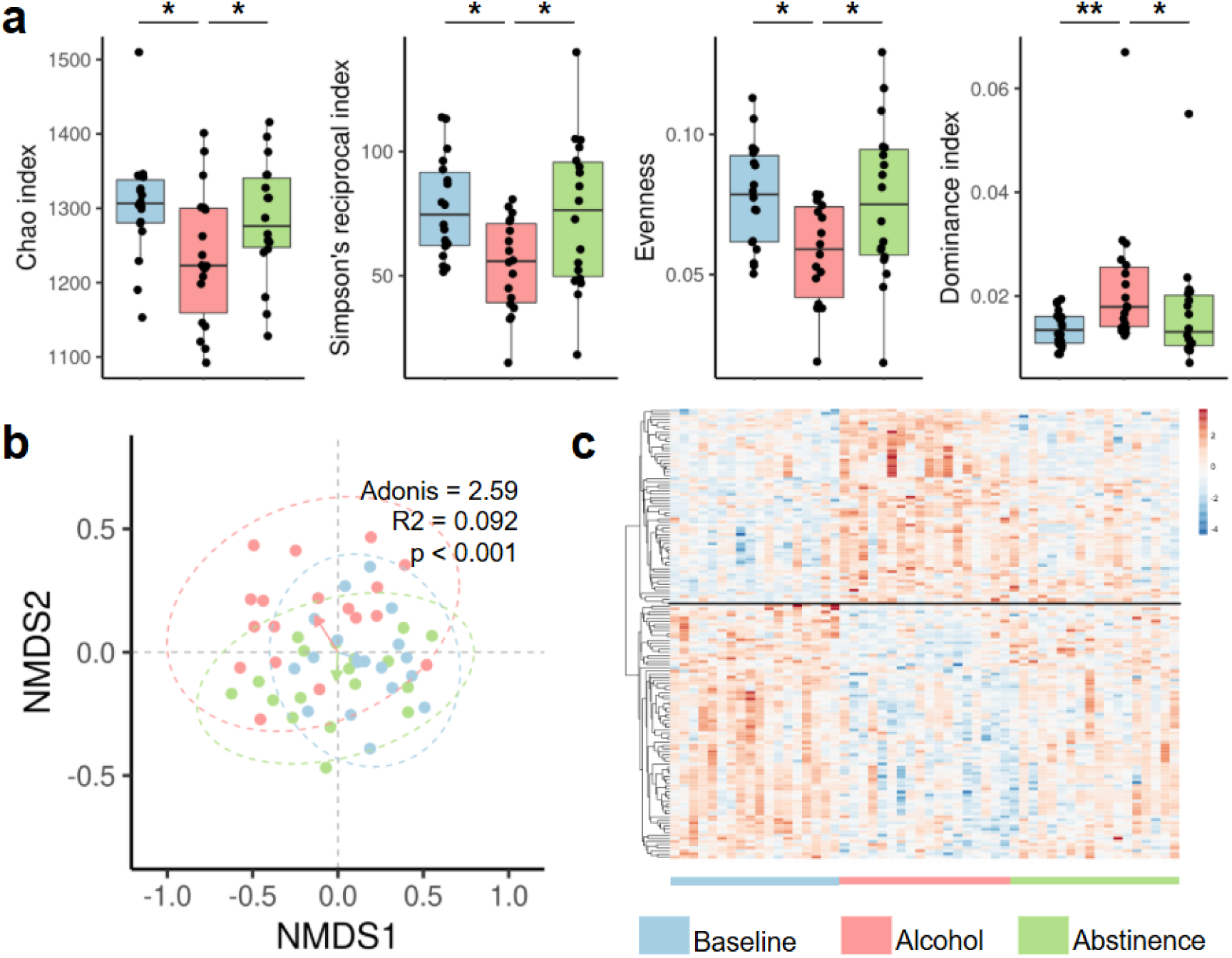
Alcohol consumption alters gut microbiota composition. (a) Alpha diversity metrics across longitudinal timepoints. Distributions were compared using pairwise t-tests or Wilcoxon rank-sum tests following assessment of normality (Shapiro–Wilk test). P-values are indicated above boxplots for Chao index (richness), Simpson reciprocal index (diversity), Simpson evenness index, and dominance index. (b) Non-metric multidimensional scaling (NMDS) of microbiota composition (vegan::metaMDS, R). Ellipses represent 95% confidence intervals, and arrows indicate centroids for each timepoint. Statistical significance of group separation (timepoint as factor) was assessed using permutation analysis (vegan::adonis2), shown as an inset. (c) Heatmap of clr-transformed abundance data for the most discriminant OTUs (n = 146). Hierarchical clustering separates alcohol-exposed samples from baseline and abstinence, indicated by the horizontal black line. Statistical significance: *p < 0.05; **p < 0.01.

To assess global compositional changes, we applied multivariate dimensionality reduction analyses, which revealed that microbial communities during baseline and abstinence were highly similar, whereas the alcohol-exposed state showed a marked divergence (Adonis = 2.59, p < 0.001). These results indicate that alcohol consumption induces pronounced microbiota alterations that rapidly revert toward baseline upon cessation (Fig. 4b).

Among the 869 OTUs, 146 showed significant differences in abundance across timepoints. Hierarchical clustering of clr-transformed abundances identified distinct clusters of taxa that were either enriched or depleted during alcohol exposure (Fig. 4c). To prioritize biologically relevant candidates, we focused on taxa with large effect sizes and clear taxonomic identification. Of the taxa enriched during alcohol exposure, only *Akkermansia* spp. (OTU29; most likely *A. muciniphila*, based on 99.75% sequence identity across the V3--V4 region) could be confidently assigned at the species level. This species exhibited an increase during alcohol exposure, which rapidly returned to baseline levels during abstinence (Supp. Fig. 4).

In contrast, several taxa formerly classified within the *Lactobacillus* genus were consistently reduced by alcohol exposure and only partially recovered during abstinence. Notably, *Limosilactobacillus* spp. (OTU286 and OTU17, related to *Lactobacillus fermentum*, and OTU459 and OTU1, related to *Lactobacillus reuteri*) showed the largest decreases associated with alcohol and remained below baseline levels after cessation (Supp. Fig. 4).

To determine whether these microbiota alterations were associated with alcohol-induced brain damage, we next integrated microbiota and neuroimaging features using a data-driven computational approach.

### Computational modeling links Akkermansia to alcohol-induced white matter alterations

Random forest models that integrated both MRI- and microbiota-derived features achieved the highest accuracy in predicting the experimental phase (baseline, alcohol exposure, abstinence). After feature selection, classification accuracy was 55.6% using microbiota features alone, 85.2% using white matter features alone, and 87.0% when both feature sets were combined (with chance level for three classes being 33.3%). These findings demonstrate that including microbiota data adds complementary predictive value to neuroimaging, improving the ability to distinguish between experimental phases.

Feature importance analyses revealed specific associations between microbiota composition and regional white matter alterations. Notably, the genus *Akkermansia* showed the strongest contribution to model performance and was consistently associated with microstructural changes across multiple tracts, including the CC and fimbria, as well as the medial forebrain bundle, cerebral peduncle, anterior commissure, and cingulum (Fig. 5).

**Figure 5.**
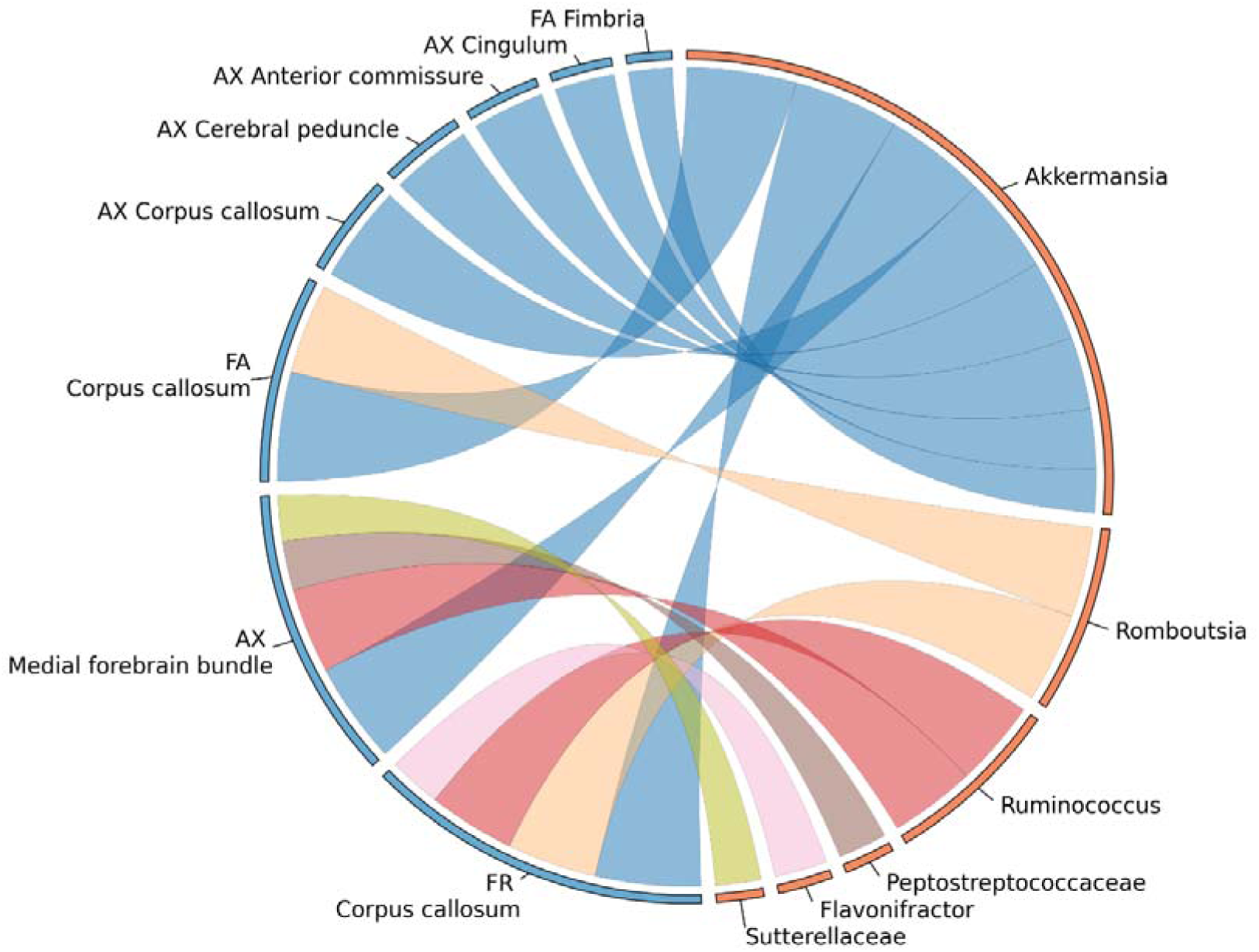
Machine learning identifies Akkermansia as a key predictor of alcohol-related white matter alterations. Circos plot illustrating associations between microbiota features (right hemicircle, orange) and MRI-derived metrics (left hemicircle, blue), based on machine learning–derived importance scores. Links represent microbiota–MRI feature pairs selected according to a combined importance score (*S_combined*), integrating three criteria: (i) the predictive power of the microbiota feature for classifying experimental phase (baseline, alcohol exposure, abstinence); (ii) the corresponding predictive performance of the MRI feature for the same task; and (iii) the ability of the microbiota feature to predict MRI feature values across phases using regression models. Link thickness is proportional to *S_combined*. Only the most informative associations are shown. FA, fractional anisotropy; FR, restricted fraction; AX, axonal radius.

Given that *Akkermansia* abundance was significantly modulated by alcohol exposure and emerged as the top predictor of white matter alterations in our integrative model, we next sought to test its functional relevance. We hypothesized that the observed expansion of *Akkermansia* during alcohol exposure might represent a compensatory response to tissue damage, and therefore evaluated whether supplementation with an *Akkermansia* strain during abstinence could mitigate alcohol-induced alterations in the gut, liver, and brain.

### Administration of Akkermansia muciniphila mitigates gut, liver, and brain pathology during abstinence following chronic alcohol exposure

Following the alcohol exposure phase, rats received either *Akkermansia muciniphila* (10 □ CFU/day) or vehicle via a gelatin matrix for six weeks during abstinence (Fig. 6a). At the end of treatment, animals were sacrificed and tissues were collected for histological and immunohistochemical analyses of the gut, liver, and brain. Both male and female msP rats were included in this experiment.

**Figure 6.**
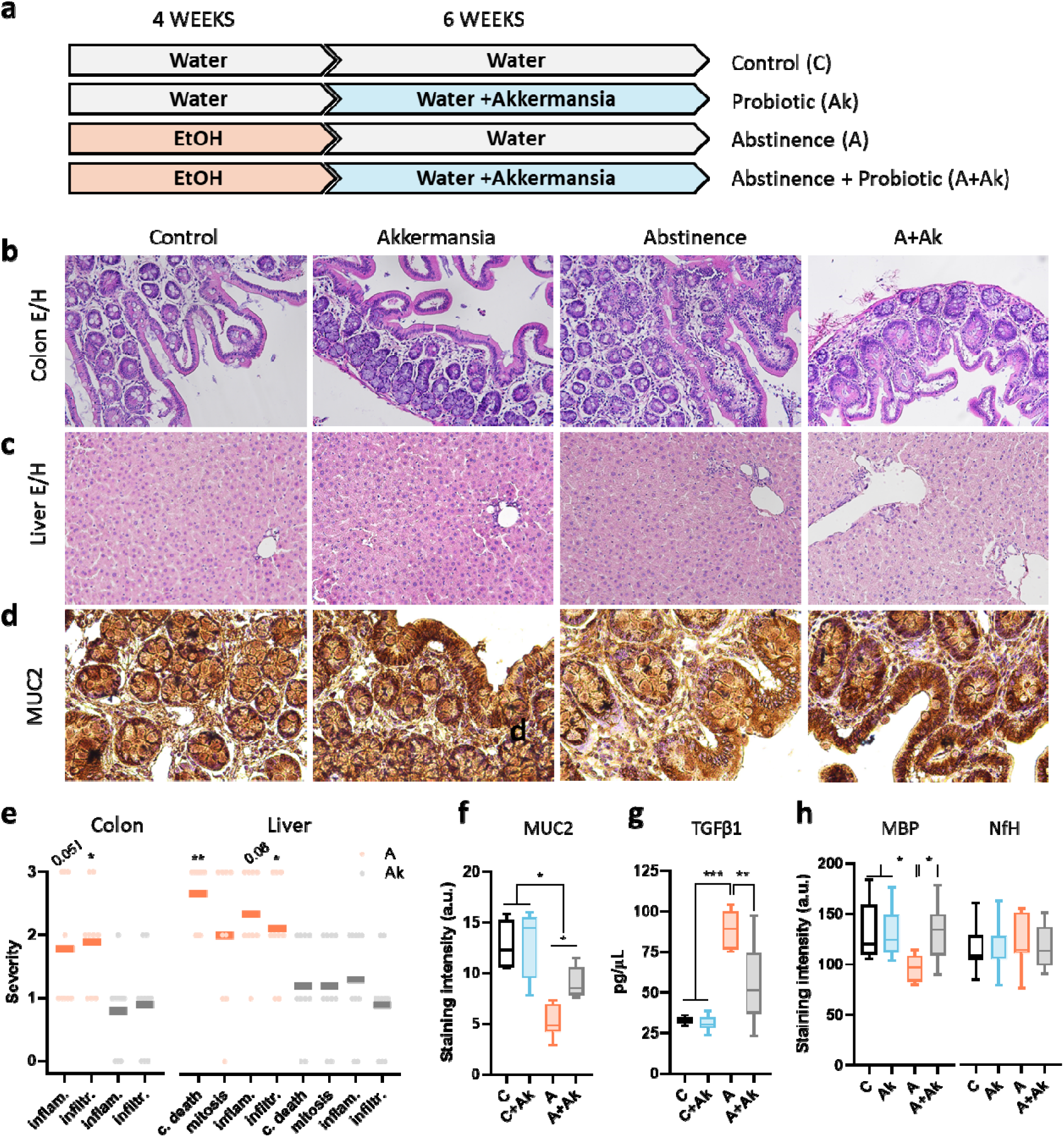
Akkermansia muciniphila supplementation mitigates alcohol-induced gut, liver, and brain alterations during abstinence. (a) Experimental design and timeline of data acquisition for the intervention study. (b–d) Representative histological images of hematoxylin and eosin (H&E) staining in colon (b) and liver (c), and mucin (MUC2) immunostaining in colon (d) across experimental groups. (e) Semi-quantitative histopathological scoring (0–3 scale; 0 = normal, 3 = severe pathology). Scoring was performed by a blinded histopathologist. (f) Quantitative analysis of colonic mucin (MUC2) levels corresponding to panel (d), shown as boxplots across experimental groups. (g) Hepatic TGF-β1 levels measured by ELISA, showing increased levels following alcohol exposure and attenuation with *A. muciniphila* supplementation. (h) Myelin basic protein (MBP) and neurofilament (NfH) immunoreactivity in the fimbria, reflecting alcohol-induced white matter alterations and their modulation by *A. muciniphila*. Statistical significance: *p < 0.05; **p < 0.01; ***p < 0.001.

We first assessed alcohol-related tissue pathology in the colon and liver using H&E staining (Fig. 6b,c) and quantified the corresponding histopathological features (Fig. 6e). Alcohol-exposed animals displayed persistent signs of gut inflammation as well as cell death, inflammation, and inflammatory infiltration in the liver, during abstinence. Notably, these alterations were significantly attenuated by *A. muciniphila* administration. In the colon, inflammation and infiltration were reduced, reaching statistical significance for infiltration (Dunn’s test: mean rank diff = 11.49, p = 0.03) and a trend for inflammation (mean rank diff = 10.57, p = 0.051). In the liver, cell death and inflammatory infiltration were significantly reduced (Dunn’s test: cell death, mean rank diff = 32.80, p = 0.003; infiltration, mean rank diff = 26.31, p = 0.027), whereas inflammation showed a trend toward reduction (mean rank diff = 22.63, p = 0.08) and mitosis remained unchanged.

To further evaluate gut barrier integrity, we quantified MUC2 levels (Fig. 6d), a major component of the intestinal mucus layer secreted by goblet cells. MUC2 was reduced in alcohol-exposed rats, indicating persistent impairment of the gut barrier during abstinence that may contribute to increased intestinal permeability (Fig. 6f). This reduction was partially reversed by

*A. muciniphila* supplementation. Two-way ANOVA revealed significant effects of drinking (F(1,23) = 45.30, p < 0.0001), probiotic treatment (F(1,23) = 5.48, p = 0.028), and their interaction (F(1,23) = 4.33, p = 0.049). Post hoc analyses confirmed a marked reduction in MUC2 levels following alcohol exposure (p < 0.0001), which was significantly attenuated by the treatment (p = 0.012).

We next assessed liver damage by measuring intrahepatic TGF-β1 levels, a marker associated with fibrosis and immune dysregulation^21^. TGF-β1 was elevated in alcohol-exposed animals and decreased toward baseline levels following *A. muciniphila* treatment (Fig. 6g). Two-way ANOVA revealed significant effects of alcohol exposure (F(1,28) = 73.41, p < 0.0001), probiotic treatment (F(1,28) = 13.46, p = 0.001), and their interaction (F(1,28) = 11.18, p = 0.0024). Post hoc analyses confirmed a marked increase in TGF-β1 following alcohol exposure (p < 0.0001) and a significant reduction with *A. muciniphila* supplementation (p = 0.005).

Finally, we examined brain white matter integrity by assessing MBP staining. As shown in Fig. 3, MBP levels were reduced after four weeks of chronic alcohol exposure, and remained low throughout the six-week abstinence period (Fig. 6h). *A. muciniphila* supplementation increased MBP in white matter to levels comparable to controls (Fig. 6h). Two-way ANOVA revealed a significant interaction between alcohol exposure and probiotic treatment (F(1,28) = 4.43, p = 0.044), with post hoc comparisons confirming a reduction in MBP following alcohol exposure (p = 0.040) and its normalization after A. muciniphila supplementation (p = 0.040).

In contrast, neurofilament staining, which was increased following alcohol exposure, had normalized by the end of abstinence and was not further modified by bacterial supplementation. Consistently, two-way ANOVA showed no significant effects of alcohol exposure, probiotic treatment, or their interaction (all p > 0.82; Fig. 6h).

Thus, alcohol cessation alone was insufficient to reverse alcohol-induced alterations in the gut, liver, and brain over six weeks of abstinence. In contrast, *A. muciniphila* supplementation during abstinence significantly attenuated intestinal and hepatic pathology and restored MBP levels in the brain to those observed in control animals, indicating recovery of myelin integrity. These findings support a beneficial effect of *A. muciniphila* supplementation on alcohol-induced gut, liver, and brain alterations and support the associations identified by our integrative model.

## DISCUSSION

In this study, we developed an integrative modeling framework that combines longitudinal diffusion MRI and gut microbiota profiling to identify microbial signatures associated with brain microstructural alterations. Applied to an established rat model of AUD, this approach showed that combining MRI and microbiota features improved prediction of alcohol exposure and abstinence states, and identified *Akkermansia* as the taxon most strongly associated with alcohol-related white matter pathology. Guided by this result, we administered *A. muciniphila* during abstinence and observed protective effects across the gut, liver, and brain. Together, these findings indicate that this framework can be used to uncover biologically relevant gut-brain relationships and to identify microbiota-based therapeutic candidates.

Analysis of brain MRI data confirmed widespread alcohol-induced abnormalities across the white matter skeleton, with particularly prominent effects in the corpus callosum, fimbria, and fornix, consistent with previous work from our group^6,7,22^. In longitudinal studies, multimodal MRI classifiers have reliably distinguished naïve, alcohol-drinking, and abstinent msP rats, and have shown sensitivity to treatment-related effects such as those induced by naltrexone^22^. Similarly, multimodal imaging features have been shown to discriminate alcohol-exposed from control brains with high accuracy and to identify disease-relevant networks in AUD patients^23^. Extending these findings, the use of advanced diffusion MRI in the present study enabled the characterization of additional microstructural features, including FR and AX, thereby increasing sensitivity to alterations in the axon–myelin unit. The combined pattern of reduced FA and FR together with altered AX indicates disruption of myelin integrity and axonal structure, a conclusion further supported by our histological analyses showing decreased MBP and altered neurofilament organization.

Longitudinal microbial profiling revealed that alcohol exposure markedly disrupted gut microbiota composition and reduced microbial diversity. These findings are consistent with reports in both animal models and patients with AUD showing reduced richness and altered community structure after chronic alcohol consumption^9,24,25^. In rodents, chronic alcohol intake has been associated with pronounced shifts in β-diversity and loss of microbial richness^23^, while in patients with AUD, microbiota disruption has been linked to increased gut permeability and systemic inflammation^24^. Such loss of diversity is thought to compromise ecosystem stability and promote inflammatory and metabolic dysfunction across the gut–liver, gut–immune, and gut–brain axes^6,26,27^. Our results extend this framework by showing that alcohol-induced dysbiosis contains information predictive of alcohol-related white matter injury.

The integrative model provided a tractable link between microbial variation and brain pathology. Within a broader set of taxa and white matter tracts, *Akkermansia* showed the strongest association with alterations in FA, FR, and AX across the corpus callosum, fimbria, cingulum, anterior commissure, cerebral peduncle, and medial forebrain bundle. These tracts include major interhemispheric and limbic pathways repeatedly implicated in AUD^28^. Other taxa, including *Romboutsia*, *Ruminococcus*, *Peptostreptococcaceae*, *Flavonifractor*, and *Sutterellaceae*, also showed robust associations, indicating that alcohol-related brain pathology is likely influenced by a consortium of microbial shifts rather than by a single taxon. Notably, several of these taxa have been linked to alcohol-induced gut dysbiosis, metabolic dysfunction, or low-grade inflammation—for example, depletion of *Ruminococcaceae* in alcohol-related liver disease (ALD), and inflammation-associated shifts in *Flavonifractor* and *Sutterellaceae* species^13,29,30^—supporting their potential contribution to systemic and brain alterations in AUD. Together, these findings support the view that multi-taxon dysbiosis converges on specific brain pathways.

A notable finding was that *Akkermansia* abundance increased during alcohol exposure and returned to baseline after cessation, which contrasts with some reports in rodents and patients with AUD where *Akkermansia* was often reduced during alcohol exposure^31–33^. For example, Addolorato et al. identified an AUD-associated microbial fingerprint characterized by reduced *Akkermansia* and increased pro-inflammatory cytokines ^32^. This discrepancy may be explained by our longitudinal design, which captured dynamic and potentially compensatory microbial responses not evident in cross-sectional studies. Furthermore, experimental evidence on the effects of alcohol on *Akkermansia* remains variable, suggesting that its response may depend on both the dose and duration of alcohol exposure^34,35^. Given the established role of *Akkermansia* in mucus turnover and barrier homeostasis^10,36^, its transient increase during alcohol exposure may reflect an adaptive response to limiting alcohol-induced damage, while the subsequent decline during abstinence could hinder gut barrier recovery and promote persistent systemic inflammation.

This interpretation is supported by the intervention study. Administration of *A. muciniphila* during abstinence partially restored gut mucosal integrity, as indicated by increased MUC2 staining, consistent with recovery of the protective mucus layer. This likely reduces epithelial exposure to luminal bacteria^37^ and limits translocation of harmful compounds to distal organs^38^. In the liver, *A. muciniphila* supplementation decreased TGF-β1 levels toward baseline and ameliorated alcohol-related histopathological changes, supporting a reduction in inflammatory and fibrotic signaling. These findings align with previous studies showing that *A. muciniphila* improves gut barrier function, increases mucus thickness, and attenuates liver inflammation in models of ALD^39,40^. Importantly, supplementation also restored MBP levels in white matter, indicating preservation of myelin-related integrity in the brain. Together, these results suggest that *A. muciniphila* supplementation has beneficial effects across multiple organs and provide experimental support for a gut-mediated contribution to persistent white matter abnormalities after alcohol exposure.

More broadly, our study illustrates how machine learning can be used not only for classification, but also for biologically informed target discovery in gut–brain axis disorders. By integrating longitudinal microbiota and neuroimaging data, this framework prioritized a candidate microorganism that subsequently showed therapeutic benefit *in vivo*. This strategy may therefore be applicable to other dysbiosis-related conditions in which peripheral microbial alterations contribute to central nervous system dysfunction. In AUD, our findings suggest that microbiota-informed intervention during abstinence may be a promising avenue to mitigate persistent gut, liver, and brain pathology.

## Data Availability Statement

All data supporting the findings of this study will be publicly available on Digital.CSIC. Further information and materials are available from the corresponding author upon reasonable request.

## Acknowledgements

Financial support for this work was provided to SC and YS by the Generalitat Valenciana and Next Generation EU through the Prometeo Excellence Grant (CIPROM/2022/15). SC further acknowledges support from the Spanish Ministerio de Ciencia e Innovación and the Agencia Estatal de Investigación (PCI2024-153491 funded by MICIU/AEI/10.13039/501100011033 and the European Union) through the ERA-NET NEURON framework project IBRAA. The authors acknowledge support from the Program for Centers of Excellence in R&D “Severo Ochoa” (CEX2021-001165-S, MICIU/AEI/10.13039/501100011033).

During the preparation of this work the authors used ChatGPT 5, from OpenAI, to improve readability and language usage. After using this tool, the authors reviewed and edited the content as needed and take full responsibility for the content of the publication.

## SUPPLEMENTARY FIGURES

**Supplementary Figure 1:**
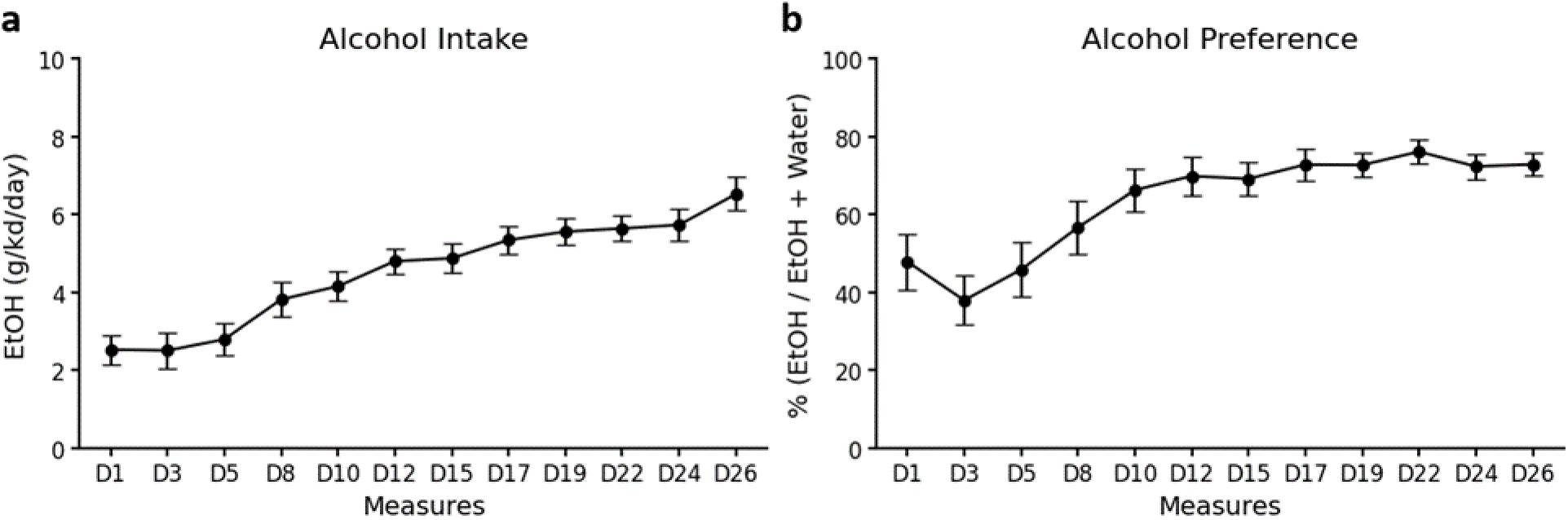
Alcohol intake and drinking preference overtime in msP rats. (a) Average alcohol intake measured in grams, adjusted for kilogram of bodyweight per day (D) for all rats. (b) Average alcohol preference measured as percentage of 10% alcohol (EtOH) consumed over total liquid consumed (EtOH + water) for all rats. Symbols represent mean ± standard error.

**Supplementary Figure 2.**
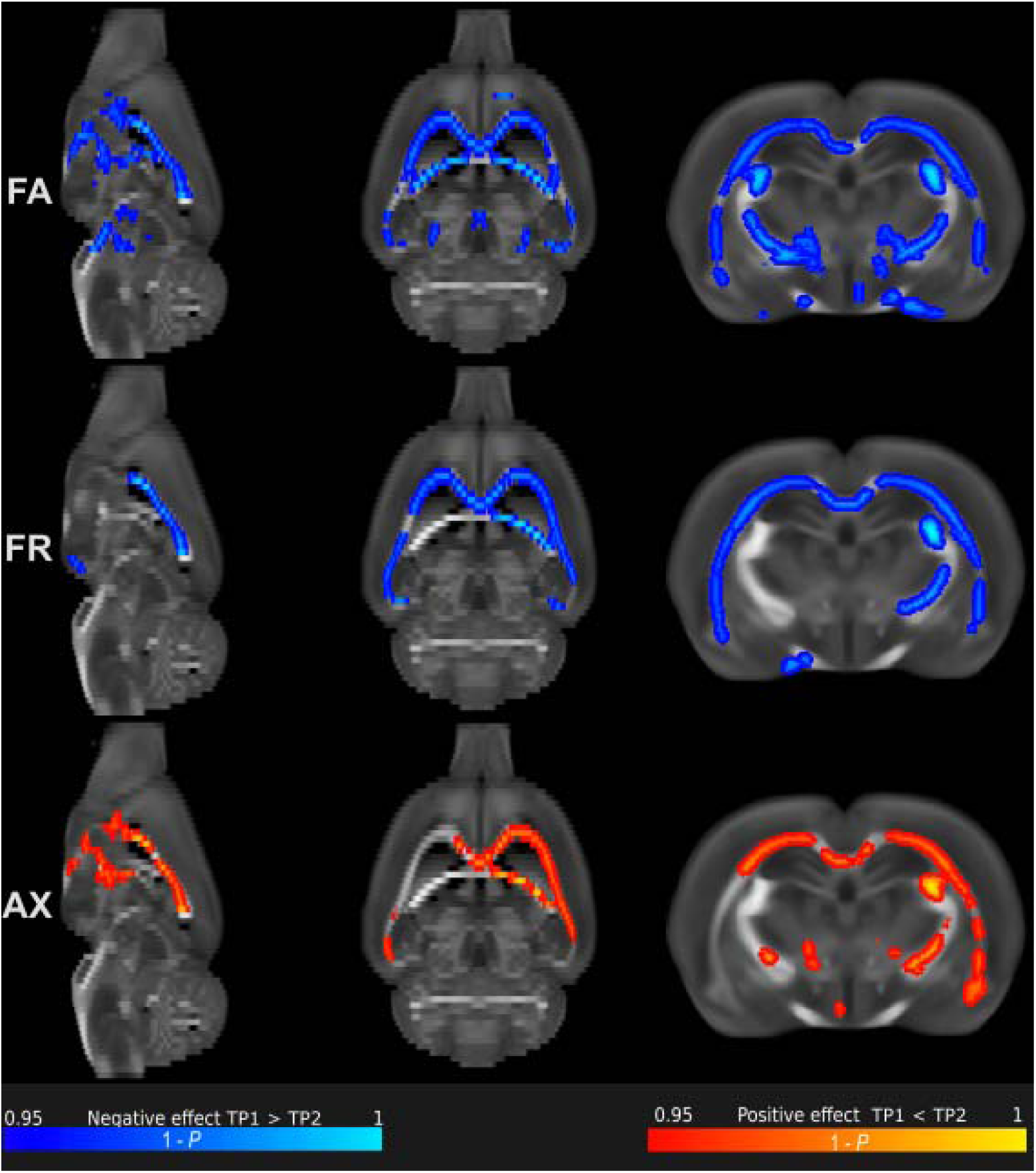
White matter alterations assessed using multiple diffusion-weighted MRI metrics. Tract-based spatial statistics (TBSS) analyses performed voxel-wise on the center of the white matter skeleton, with threshold-free cluster enhancement correction, revealed widespread longitudinal microstructural changes in the alcohol-exposed group. Specifically, reductions in fractional anisotropy (FA) and restricted fraction (FR) were observed from baseline to the alcohol-drinking condition, accompanied by an increase in axonal diameter (AX).

**Supplementary Figure 3:**
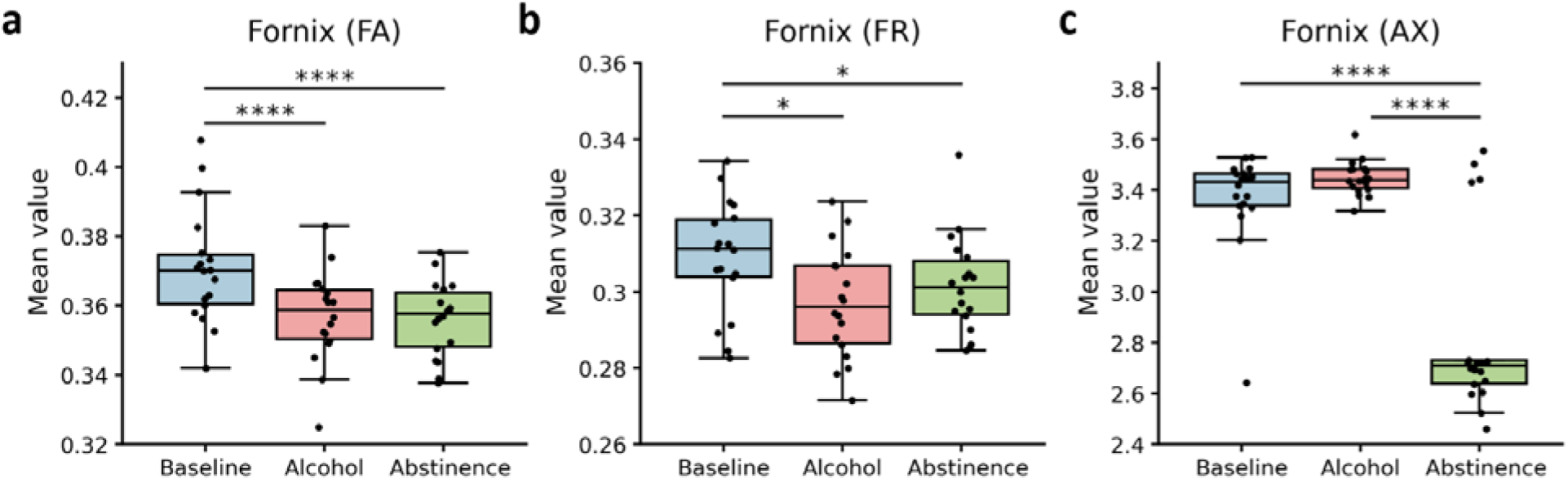
Dw-MRI metrics in the Fornix. Boxplots representing MRI-derived measurements of: a) Fractional Anisotropy (FA), b) Restricted Fraction (FR), and c) Axonal Diameter (AX) for the Fornix in each phase of alcohol consumption (baseline, alcohol, abstinence). Box size denotes the 25^th^ and 75^th^ percentiles, with the line showing the median. Whiskers extend 1.5 times the interquartile range from the 25^th^ and 75^th^ percentiles. Statistical significance: *p*<0.05 (∗), *p*<0.01 (∗∗), *p*<0.001 (∗∗∗), p<0.0001 (∗∗∗∗). Individual rats are represented by dots.

**Supplementary Figure 4:**
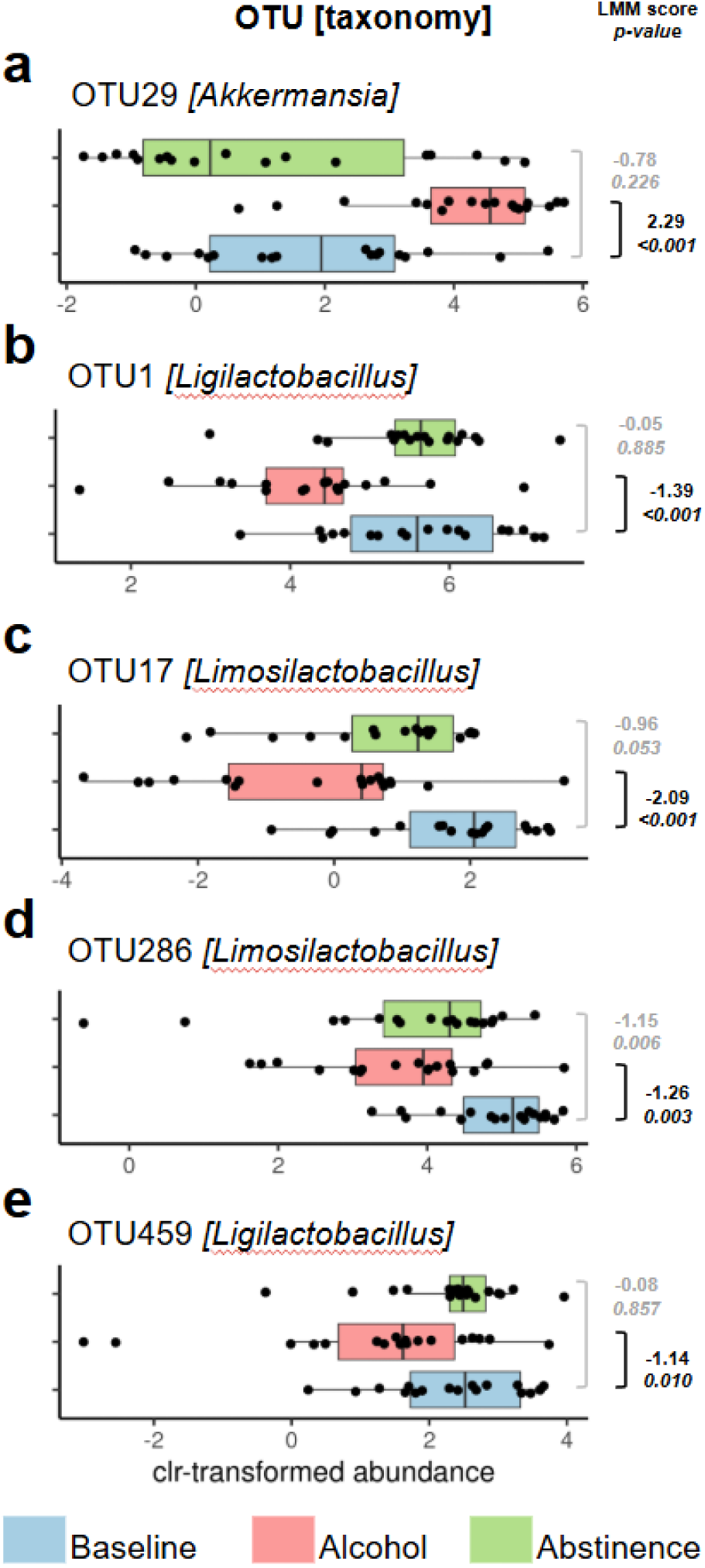
Alcohol consumption causes shifts in the rat gut microbiota. (a-e) Differential abundance for selected OTU features is shown as boxplots. OTUs were selected according to extreme statistical estimates resulting from applying linear mixed models (LMM, nlme::lme R function) and assuming idiosyncratic variation of animal specimens as a random variable and reaching certain taxonomy assignment at least at the genus level. The respective statistical estimates and p-values are shown for every pairwise comparison. Taxonomy identification was assisted using SINA aligner web tool and SILVA 138 nomenclature.

**Supplementary Figure 5:**
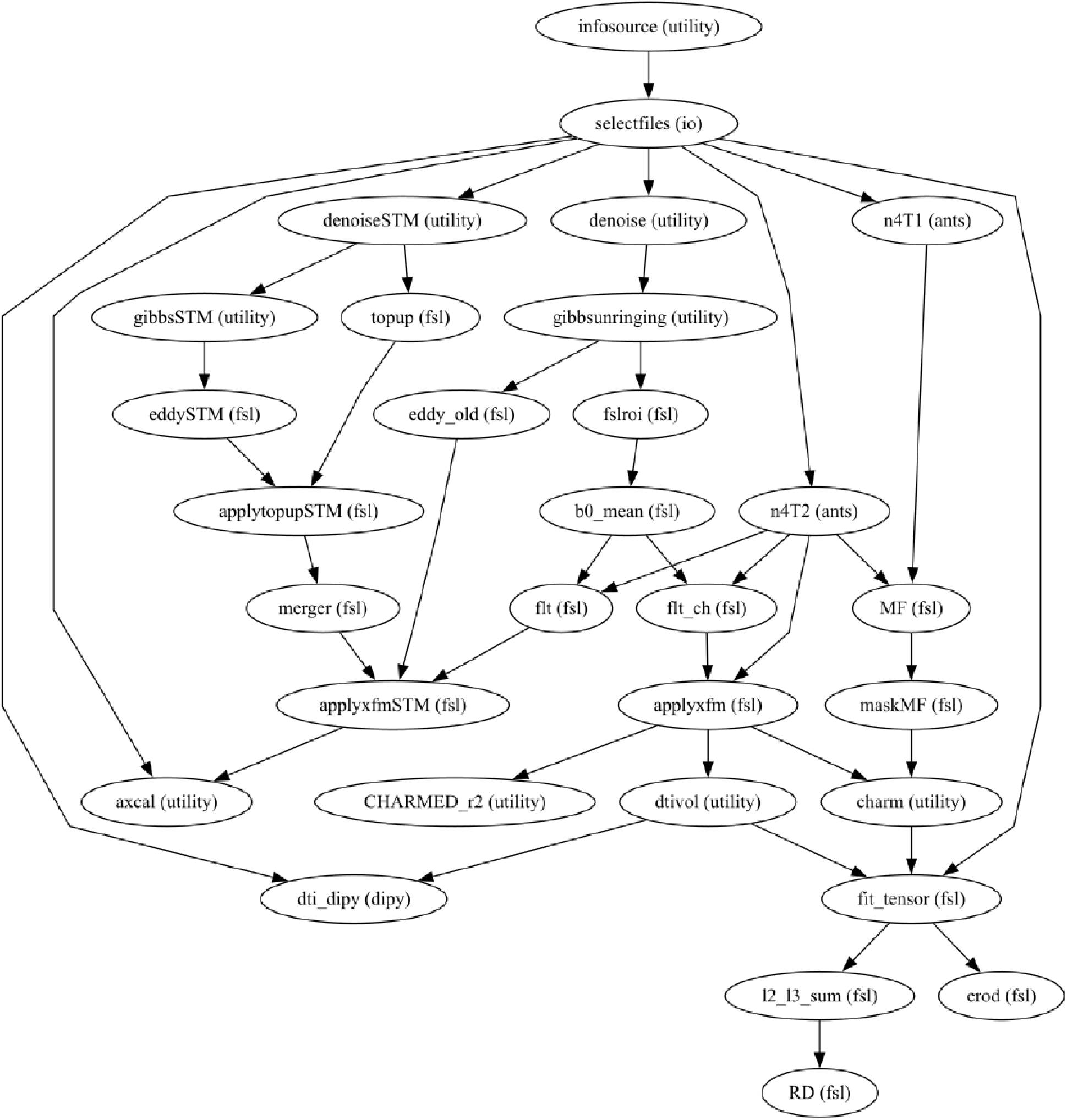
Preclinical data preprocessing pipeline. Diffusion-weighted MRI data preprocessing steps including denoising, motion correction and model fitting.

## SUPPLEMENTARY TABLES

**Supplementary Table 1.**
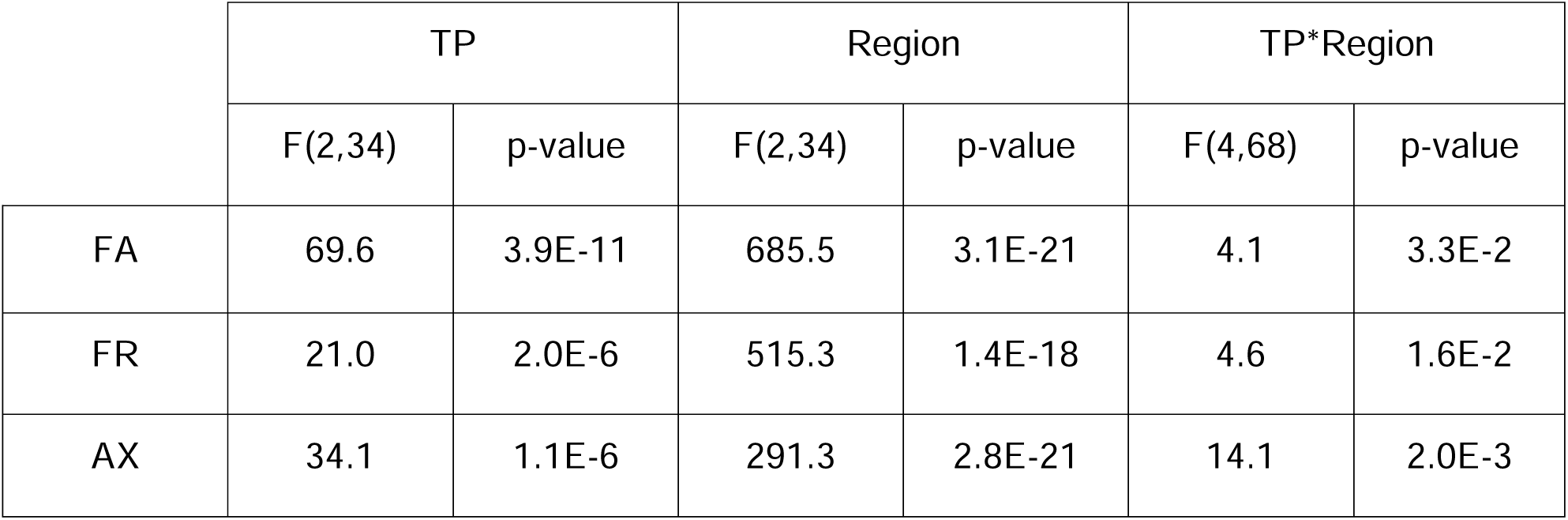
Two-way ANOVA results for diffusion MRI metrics across timepoints (TP: baseline, alcohol exposure, and abstinence) and regions of interest (Region: corpus callosum, fimbria, and fornix) in the experimental group. Significant main effects of timepoint and region were observed for all metrics (FA, fractional anisotropy; FR, restricted fraction; AX, axonal radius), along with a significant timepoint × region interaction, indicating region-specific longitudinal microstructural changes. F values and corresponding p-values are reported for each factor and interaction.

|  | TP |  | Region |  | TP*Region |  |
| --- | --- | --- | --- | --- | --- | --- |
|  | F(2,34) | p-value | F(2,34) | p-value | F(4,68) | p-value |
| FA | 69.6 | 3.9E-11 | 685.5 | 3.1E-21 | 4.1 | 3.3E-2 |
| FR | 21.0 | 2.0E-6 | 515.3 | 1.4E-18 | 4.6 | 1.6E-2 |
| AX | 34.1 | 1.1E-6 | 291.3 | 2.8E-21 | 14.1 | 2.0E-3 |

**Supplementary Table 2.**
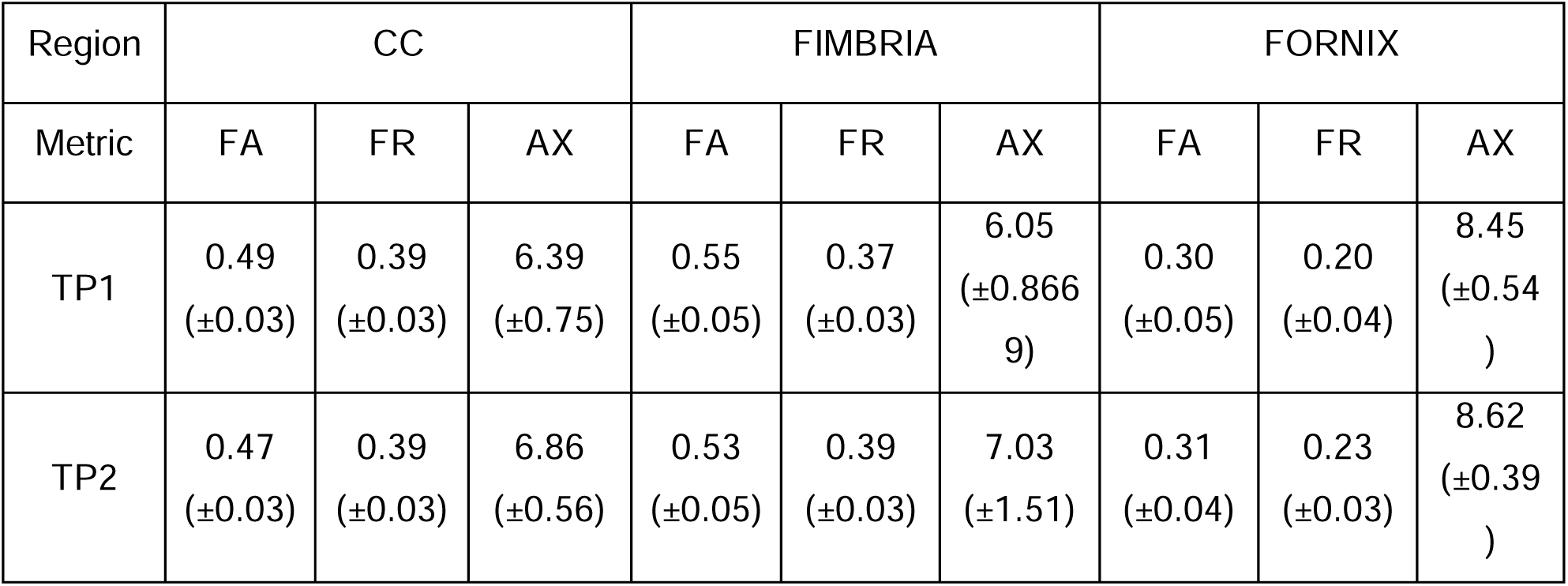
Mean (± SD) values of the three MRI metrics across regions of interest and timepoints in the control (water-drinking) group. Two-way ANOVA showed no significant effects for FA or FR, and a significant main effect of timepoint for AX (F(1,8) = 8.1, p = 0.02). However, post hoc analyses showed no significant differences across regions, with a trend toward change in the fimbria (CC: t (8) = −1.6, p = 0.16; fimbria: t (= 8) = −2.0, p = 0.09; fornix: t (8) = −1.0, p = 0.34).

